# Noise regularization removes correlation artifacts in single-cell RNA-seq data preprocessing

**DOI:** 10.1101/2020.07.29.227546

**Authors:** Ruoyu Zhang, Gurinder S. Atwal, Wei Keat Lim

**Affiliations:** Regeneron Pharmaceuticals Inc., Tarrytown, New York

## Abstract

With the rapid advancement of single-cell RNA-seq (scRNA-seq) technology, many data preprocessing methods have been proposed to address numerous systematic errors and technical variabilities inherent in this technology. While these methods have been demonstrated to be effective in recovering individual gene expression, the suitability to the inference of gene-gene associations and subsequent gene networks reconstruction have not been systemically investigated. In this study, we benchmarked five representative scRNA-seq normalization/imputation methods on human cell atlas bone marrow data with respect to their impact on inferred gene-gene associations. Our results suggested that a considerable amount of spurious correlations was introduced during the data preprocessing steps due to over-smoothing of the raw data. We proposed a model-agnostic noise regularization method that can effectively eliminate the correlation artifacts. The noise regularized gene-gene correlations were further used to reconstruct gene co-expression network and successfully revealed several known immune cell modules.

## Introduction

Gene co-expression network analysis is a common approach to gather biological information and uncover molecular mechanisms of biological processes. Microarray and RNA-seq data of bulk cells have been successfully used to infer gene-gene correlations and further reconstruct gene co-expression networks^1,2^. However, these approaches are limited to measuring average gene expression across pool of mixed cell types. Single-cell RNA-seq (scRNA-seq) technology makes it possible to profile gene expression at single cell resolution, which allows for dissection of the heterogeneity within the superficially homogenous cell populations and identification of hidden gene-gene correlations masked by bulk expression profiles^3,4^.

The rapid development of scRNA-seq technology provides the opportunity to gain new insights into complex biological systems. However, due to various factors in single-cell experiments, such as differences in cell lysis, reverse transcription efficiency, and molecular sampling during sequencing^5^, scRNA-seq data are generally highly variable and noisy. To address these issues, numerous data preprocessing methods have been proposed for scRNA-seq data analysis, which generally fall into two major categories: (i) transcript abundance normalization, and (ii) dropout imputation. The observed sequencing depth can vary dramatically from cell to cell. Data normalization is hence required to remove the technical noise while preserving true biological signals. scRNA-seq data is further complicated by high dropout rate^6,7^, which refers to the phenomenon by which a large proportion of genes have a measured read count of zero due to technical limitation in detecting the transcripts rather than true absence of the gene. Data imputation has been proposed to handle the dropouts and recover the undetected gene expressions.

scRNA-seq data preprocessing methods have been benchmarked for various tasks, such as cell clustering, detection of differentially expressed genes and trajectory analysis^8^. The suitability of these methods for reverse engineering gene networks and, in general, for measuring gene-gene association, has not been systemically evaluated. Andrews *et. al* tested several imputation methods on a small simulation dataset and found that dropout imputation would generate false positive gene-gene correlations^9^. However, the simulation dataset in that study represented the simplest case without technical confounders, thus the effect of data preprocessing on real data remains unknown.

In this study, we benchmarked five normalization/imputation methods, which are representatives of their own methodology groups, in respect to their influence on gene-gene correlation inferences. The first method, global scaling normalization, normalizes a cell’s gene expression levels (usually measured by the unique molecule index (UMI)) by its summed expression over all genes, e.g. total UMI. This method is usually followed by log transformation and z-score scaling in the downstream analyses. Since the log transformation and z-score scaling are monotonic (rank-preserved) functions, we only included total UMI normalization in our benchmarking (referred as NormUMI). The second normalization framework utilizes “Regularized Negative Binomial Regression” to normalize and stabilize variance of scRNA-seq data (referred as NBR). This method showed remarkable performance in removing the influence of technical noise while preserving biological heterogeneity^10^. Three imputation methods were also included: (i) MAGIC, a data smoothing approach which leverages the shared information across similar cells to denoise and fill in dropout values^11^, (ii) SAVER, a model based approach which models the expression of each gene under a negative binomial distribution assumption and outputs the posterior distribution of the true expression^12^, (iii) DCA, an adapted autoencoder framework which is able to capture the complexity and non-linearity in scRNA-seq data and infer gene expressions^13^.

To evaluate the influence of these preprocessing methods on gene-gene correlation inference, we applied them to bone marrow scRNA-seq data from Human Cell Atlas Project^14^. We computed gene-gene correlation after the data preprocessing and compared results among the methods. With the exception of NormUMI, the normalization method with the least data manipulation, all other normalization/imputation methods presented a noticeable inflation of gene-gene correlation coefficients and introduced correlation artifacts for gene pairs that are not expected to be coexpressed. In addition, gene pairs with the highest correlations inferred from these methods had weak enrichments in protein-protein interactions from STRING database^15^, suggesting that many of these correlations may be the false signals introduced during the data preprocessing. Further data inspection using random and non-associated gene pairs as negative control indicated that the artifacts could be generated from data over-smoothing. In machine learning, adding noise under certain conditions has previously been shown to increase robustness of the results and reduce overfitting^16–18^. To this end, we implemented a noise regularization step to the preprocessed scRNA-seq data by adding uniformly distributed noise that is scaled to the dynamic expression range of each individual gene. We found that this additional step efficiently reduced gene-gene correlation artifacts and improved overall evaluation metrics. We used the regularized expression data to reconstruct gene co-expression network and successfully revealed several known immune cell modules. The canonical cell type marker genes were also rated higher in network topological properties, e.g. degree and PageRank, pinpointing their key role in their respective cell clusters.

## RESULTS

### Compute gene-gene correlation using scRNA-seq data

Previous benchmarking studies on scRNA-seq data preprocessing methods were mostly based on simulated datasets with certain assumptions in the simulation process that might not be representative of real-world data. Depending on the simulation algorithm used, results might be biased toward certain methods. For instance, the method SAVER, which uses negative binomial distribution to model and impute the data, will stand out if the simulated dataset is also generated based on a negative binomial model. To avoid such biases, we employed real world bone marrow scRNA-seq data from Human Cell Atlas (HCA) Preview Datasets as our benchmarking dataset^14^ for various data preprocessing methods. The full dataset contains 378,000 bone marrow cells which can be grouped into 21 cell clusters (**Fig. S1**), covering all major immune cell types. We randomly sampled 50,000 cells from the original dataset and excluded genes expressing in less than 100 cells (0.2%) in this subset. The final benchmarking dataset contains 12,600 genes that could form over 79 million possible gene pairs.

Five representative data preprocessing methods were applied to the single-cell expression data matrix, including two normalization methods NormUMI, NBR, and three imputation methods DCA, MAGIC and SAVER (**Fig. 1**). An important merit of scRNA-seq is its ability to unbiasedly capture the whole transcriptome of different cell types in a heterogenous cell population. Expression of two genes could be highly correlated only in one specific cell type and therefore revealing cell type-specific gene-gene associations. To capture the correlations across different cell types, we computed spearman correlation of gene pairs within the 10 largest clusters (>500 cells per cluster) in our benchmarking dataset, which included CD4 T cell, CD8 T cell, natural killer cell, B cell, Pre-B cell, CD14+ Monocytes, FCGR3A+ Monocytes, Erythrocytes, Granulocyte-macrophage progenitors and Hematopoietic stem cells (**Fig. 1**, **Fig. S1**). The highest correlation among these 10 clusters were recorded as the final correlation for each gene pairs (**Methods**).

**Figure 1.**
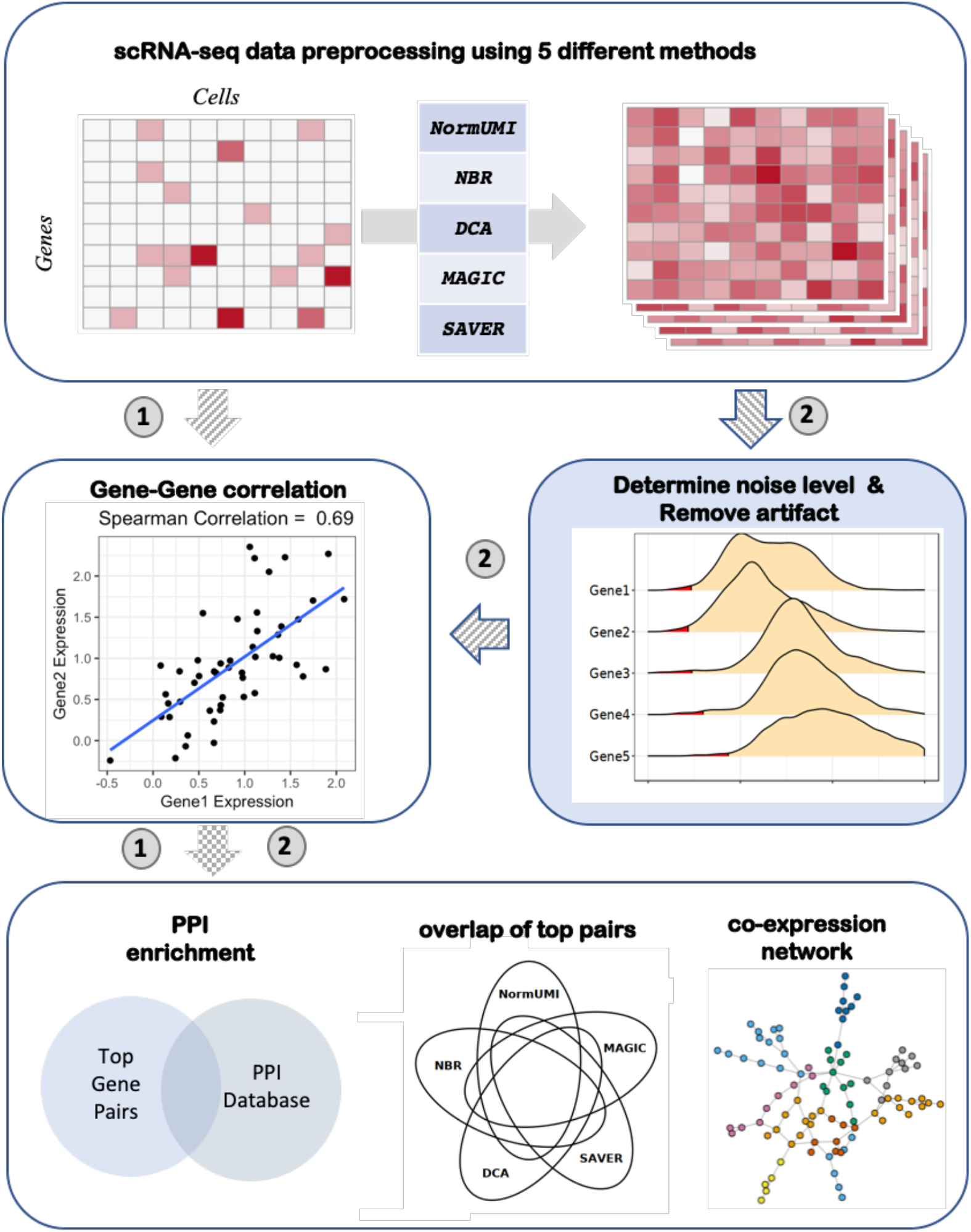
Overview of the benchmarking framework. Five scRNA-seq data preprocessing methods were applied to bone marrow single-cell expression data matrices. The gene-gene correlations were first calculated directly from the matrices after data preprocessing (denoted as route 1). We evaluated the methods by their derived gene-gene correlations enrichments in STRING PPI database as well as the consistency between methods. The evaluation results indicated that the data preprocessing procedure introduced artificial correlations. We then introduced a noise regularization step (denoted as route 2): random noise generated based on gene expression level (red areas) was applied to the expression matrices before proceeding to correlation calculation. This noise regularization step effectively reduced the spurious correlations, and the refined gene-gene correlations can be used to construct gene co-expression networks.

### Data pre-processing introduced spurious correlations

We first compared the distribution of the overall gene-gene correlations calculated from data matrices processed by the five methods. Since most of the gene pairs are not expected to have any association, we anticipate that the correlation distributions should peak around 0. However, with the exception of NormUMI, all other methods produced much higher median correlations values (NormUMI ρ=0.023, NBR ρ=0.839, MAGIC ρ=0.789, DCA ρ=0.770, SAVER ρ=0.166) (**Fig. 2A**). We proceeded to assess whether a higher correlation, after a specific data preprocessing method, would reflect a higher chance of either functional or physical interaction between the two genes. Proteins encoded by a co-expressed gene pair are more frequently interacting with each other than a random pair. Therefore, if the resulting higher correlations are true positives, they should have relatively higher enrichment in protein-protein interactions (PPI) database, while the spurious correlations would dilute the enrichment. We used the STRING database^15^, which contains 5,772,157 interacting gene pairs, to evaluate the PPI enrichment of the top correlated gene pairs derived from each method. We selected top gene pairs (ranked by correlation coefficients) from each method and calculated the overlapping fraction of these pairs with the STRING database (**Fig. 2B**). Our results showed that NormUMI had the highest PPI enrichments: 80% and 47% overlapped with STRING in the top 100 and 10,000 gene pairs, respectively. On the contrary, the top gene pairs from NBR had very low overlap with STRING (<2%), while MAGIC and DCA had similar PPI enrichments, ranging from 11% to 22%. SAVER yielded relatively better results, but the enrichments were merely half of those by NormUMI. We also randomly sampled gene pairs and overlapped the random pairs with PPI to estimate the background enrichment level (**Fig S2**). The estimated background enrichment level was ~3.6%, indicating that PPI enrichment of NBR was even lower than the background. Although this is a rather naïve method that directly relates physical interactions with gene coexpression, the results here should still provide a fair comparison among the data preprocessing methods given that the same assumption is made for all of them.

**Figure 2.**
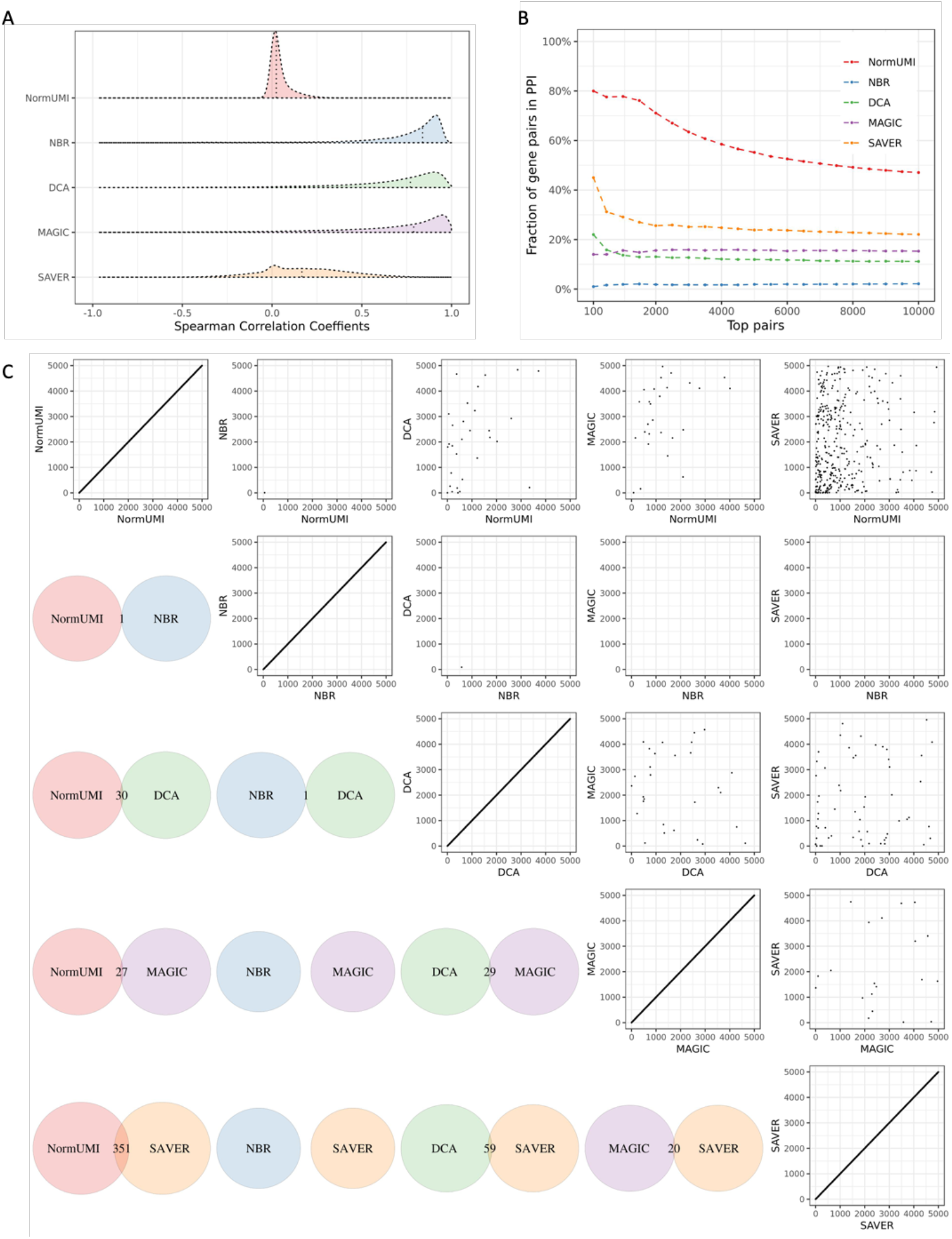
Spurious gene-gene correlations are introduced during data preprocessing. (A) The distributions of the calculated correlations varied by preprocessing methods. NormUMI had a distribution centered close to zero, while NBR, DCA and MAGIC all had apparently inflated correlation distributions. Vertical dotted lines indicate correlation medians. (B) Enrichment curves of the top correlated gene pairs in PPI for each method. X axis indicates the top n gene pairs ranked by spearman correlation coefficients; y axis indicates the fraction of the n genes pairs appearing in the STRING PPI database. NormUMI had the highest enrichment, followed by SAVER, MAGIC, DCA and NBR. (C) There was low consistency between the methods in inferring highly correlated gene pairs. Lower triangle indicates the overlapping of the top 5000 gene pairs between the two denoted methods. The largest overlap was between NormUMI and SAVER, which has only 351 (~7%) gene pairs ranked in the top 5000 in both methods. Upper triangle compares the exact rank of the shared gene pairs between methods, which also shows low levels of agreement.

Bona fide gene-gene co-expression should be identified regardless of the data preprocessing methods. To test this, we compared the consistency of highly correlated gene pairs derived from the five data preprocessing procedures. We did a pairwise comparison of the top 5,000 gene pairs selected from each method and found that the overlapping gene pairs among methods was minimal. Only 1 gene pair was shared between NormUMI and NBR out of the top 5,000 pairs. The highest overlap was between NormUMI and SAVER, with only 351 pairs (~7%) shared by the two methods (lower triangle in **Fig. 2C**). We further compared the ranks of the shared pairs between the methods and found that there was also no clear trend in their top inference (upper triangle in **Fig. 2C**). While this is not a fully quantitative assessment, it is clear that the high correlations derived from these data preprocessing methods are likely to be artifacts.

### Negative control

We next inspected several “negative control gene pairs” to get some insights into the potential cause of the spurious correlations. We defined a negative control pair using the following criteria: the two genes should not (i) appear as an interacting pair in STRING database; (ii) share any gene ontology (GO) term^19,20^; (iii) be on the same chromosome. As an example, one of the negative control gene pairs, MB21D1 and OGT, had high correlation after data processing by NBR (ρ=0.843), DCA (ρ=0.828) and MAGIC (ρ=0.739) in cell cluster #2. We also calculated the mutual information (MI) of the negative gene pairs, which can assess the strength of the association between two variables even when the relationship is highly nonlinear^21^. In this negative pair example, NBR (MI=2.10 nat), DCA (MI=0.72 nat) and MAGIC (MI=0.663 nat) also showed much higher mutual information than the other two methods, NormUMI (MI=5×10^−5^ nat) and SAVER (MI=0.053 nat). Scatter plots of the gene pair expression values after data preprocessing are shown in **Fig. 3**. Out of the five methods, NormUMI was the only method that retains the zero counts from the raw data. From NormUMI, 6110 cells out of 6534 cells (93.5%) had zero values in both genes, 3 (0.04%) cells had non-zero values in both genes, while 1.3% and 5.2% cells had non-zero for MB21D1 and OGT, respectively. The other imputation methods intensely altered the zeros from the original expression matrix. We observed that after these procedures, the processed data all presented some degree of over-smoothing, especially in the “double zeros regions” in the original data, which created the correlation artifacts (**Fig. 3**). Although NBR was not an imputation method and only shifted the zero values minimally, artificial rank correlations were introduced due to the difference in the adjusted magnitude per cell.

**Figure 3.**
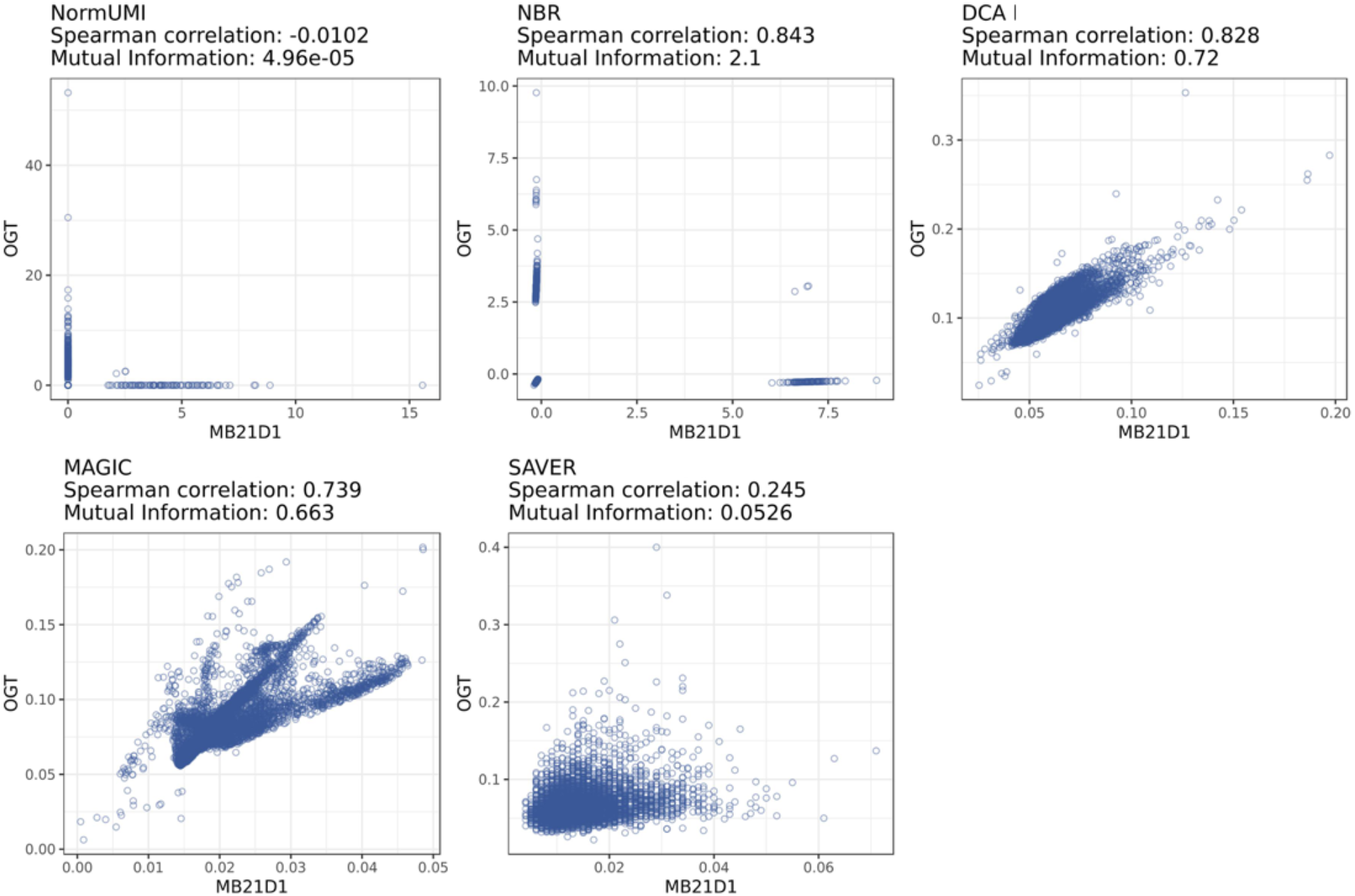
Spurious gene-gene correlation caused by data over-smoothing. Scatter plot of expression values of non-associated gene pair, OGT and MB21D1, preprocessed by different methods. There is no existing evidence to indicate that these two genes are correlated and only 3 out of 6534 cells in cluster #2 had non-zero expression value in both genes in the original expression matrix. However, after preprocessing, NBR, DCA and MAGIC all produced high correlations (0.843, 0.828 and 0.739) and high mutual information (2.1, 0.72 and 0.663 nat) between these two genes. The visualization suggested this correlation artifact may be caused by data over-smoothing.

### Noise regularization reduced the spurious correlations

Regularization is a commonly used approach to prevent overfitting/over-smoothing in machine learning and a previous work has demonstrated an equivalent form of regularization by introducing noise^16^. Here, we proposed a method utilizing noise to penalize oversmoothed expression data and further reduce spurious correlations. To implement the method, we added random noise to every single feature in the expression matrix processed by the above preprocessing methods. Taking the expression value of gene *i* in cell *j*, denoted as *V*, as an example, we generated the noise by the following steps: (i) calculate the expression distribution of gene *i* after data preprocessing procedure; (ii) determine the 1 percentile of expression value of gene *i*, termed *M*, to be used as the maximum of noise level (**Fig. 1**); (iii) generate a uniformly distributed random value, ranging from 0 to *M*, and add this random value to *V*.

After applying noise regularization to the data matrices produced by each preprocessing method, we recomputed the gene-gene correlations. The correlation medians shifted towards 0 for all five methods (**Fig. 4A**), indicating a reduction in the correlation inflation. There were also substantial improvements in the PPI enrichment for all methods (**Fig. 4B**). NBR, which previously had the lowest enrichment, yielded the highest PPI enrichment after noise regularization. In the top 100, 1,000, and 10,000 gene pairs in NBR, 99.0%, 96.8% and 67.7% could be found in the PPI database, corresponding to 99.0-, 50.9- and 31.6-fold improvement, respectively. DCA on average had ~12% PPI enrichment in previous results. After noise regularization, it produced 97.6% enrichment in the top 100 pairs and 55.8% in the top 10,000 pairs, corresponding to a ~5-fold improvement. NormUMI, which had the highest enrichment before noise regularization, also benefitted from a ~1.1 to 1.3-fold improvement. To test the robustness and reproducibility of the noise regularization results, we repeated the procedure 10 times with different random seeds to generate random noise and observed that the PPI enrichment performance were stable between repeats. The standard deviation of NBR in most points were less than 0.1% (error bar represents 99% confidence interval in **Fig. 4B**).

**Figure 4.**
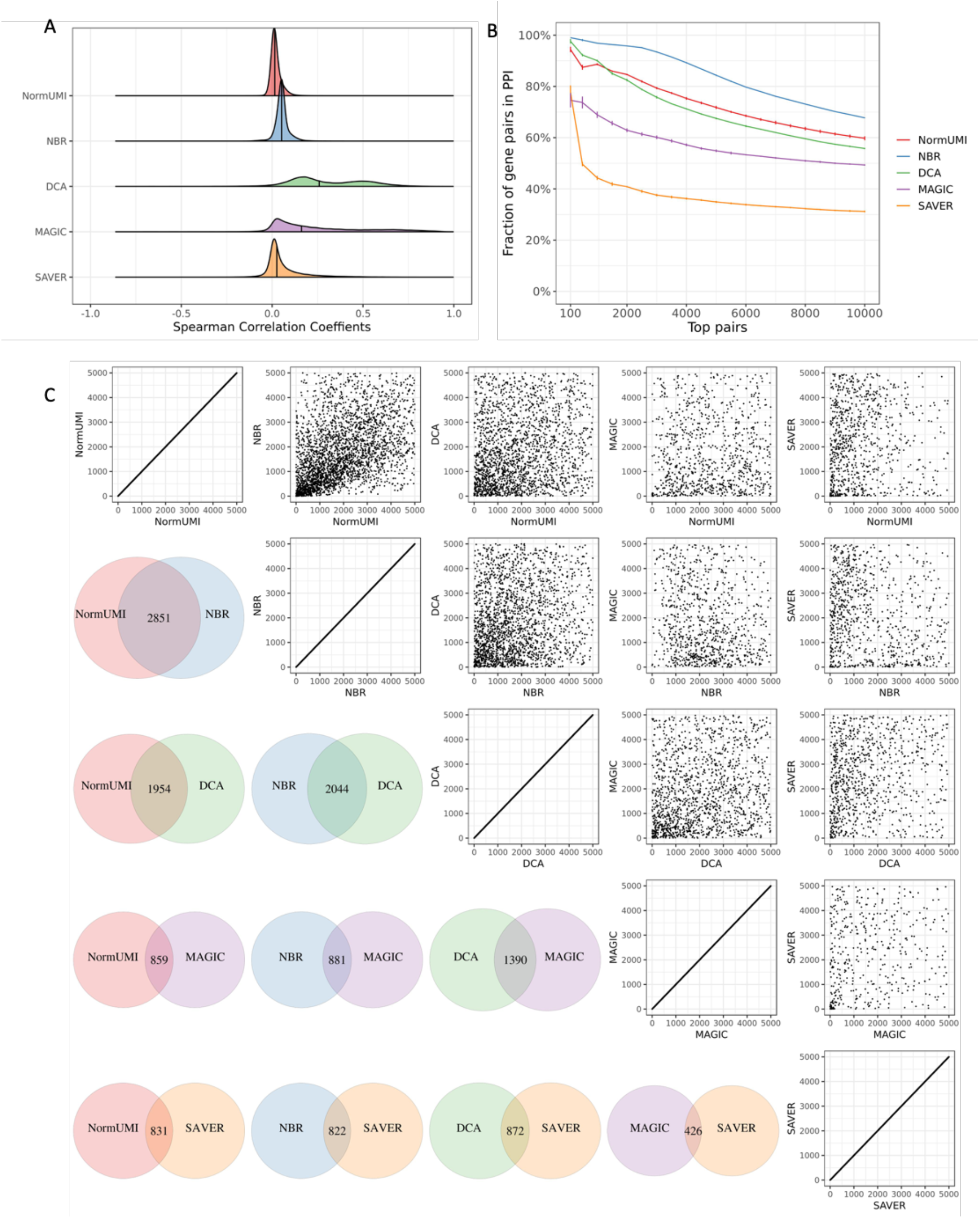
Noise regularization reduces spurious correlations. (A) After applying noise regularization, previously inflated correlation distributions from each method shifted toward zero. Vertical solid lines indicate correlation medians. (B) There were substantial improvements of the PPI enrichment in the top correlated genes. Error bars indicate 99% confidence interval based on 10 replicates, assuming error follows a Gaussian distribution. (C) Comparing to previous un-regularized data (**Fig. 2C**), there are higher levels of agreement among different methods. For example, more than 50% gene pairs were shared between NormUMI and NBR.

Different methods also showed higher agreements after applying noise regularization. Among the top 5000 gene pairs, there were 2851 (57%) overlapped between NormUMI and NBR (**Fig. 4C** lower triangle) and there was a significant correlation between the overlapped gene pairs (spearman correlation, ρ=0.50; Fisher’s exact test, *p*=1.77×10^−181^, **Fig. 4C** upper triangle). We also observed a higher degree of commonly identified gene-gene correlations between the other preprocessing methods, particularly between the top gene pairs.

### Gene-gene correlation network inferred from single-cell RNA-seq data

Co-expression networks can be used to identify gene modules with common biological functions, upstream regulators, and physically interacting proteins^22^. With the gene expression measurement at single cell resolution, scRNA-seq has fostered discoveries by improving our understanding of biological processes under different cell contexts. Therefore, gene-gene correlations revealed from single cell also have the potential to reconstruct more comprehensive networks uncovering cell type specific modules. Here, we used gene-gene correlations derived from NBR with noise regularization, since it yields the highest PPI enrichment among all the methods. In order to focus more on cell type-specific interactions, we removed housekeeping genes that typically reflect the general cellular functions and are expected to express in all cells regardless of cell type. There were 3,984 house-keeping genes removed from the original 12,600 genes. The 1000 gene pairs with the highest correlations were then taken from each cluster (cluster #0 to cluster #9) to reconstruct the network. Degree and PageRank, two algorithms from graph theory, were used to measure the importance of each gene in the network. The degree of a gene in a network is simply the number of links (interactions) the gene has^23^. Important genes tend to connect with many other genes and therefore should have relatively high degrees. In addition to the quantity of links, PageRank also takes into consideration quality of links to a gene and measures the overall ‘popularity’ of a gene^24^.

We compared the gene co-expression networks reconstructed from pre- and post-regularized data. Results show that the latter network better represented the biological functions in the topological structure and has higher degree or PageRank genes with more important functions in the immune system. For instance, LYZ, CD79B and NKG7, the canonical marker genes for monocytes, B cells and natural killer cells, respectively, yield higher PageRank and degree in the network with noise regularization. On the contrary, CD79B and NKG7 do not exist at all in the network without noise regularization (**Fig. 5 A** and **B**). We next overlaid existing PPI evidence to further refine the network by retaining only gene pairs from the STRING database^25,26^. An algorithm providing efficient visualization of different network modules, EntOptLayout^27^, was applied and the network revealed several cell type related modules that can be associated with the known biology in our benchmarking dataset (**Fig. 5C**). For instance, the upper right corner represents the B cell and Pre-B cell module, with CD79A and CD79B having higher PageRank values that are proportional to the node size. Similarly, the natural killer cell module is represented in the lower right corner, and the middle right section represents T cell as well as a transit from cytotoxic CD8 T cell to natural killer cell. These results demonstrate that, after implementing noise regularization, scRNA-seq data can be used to reconstruct gene coexpression networks that better reflect the underlying biology.

**Figure 5.**
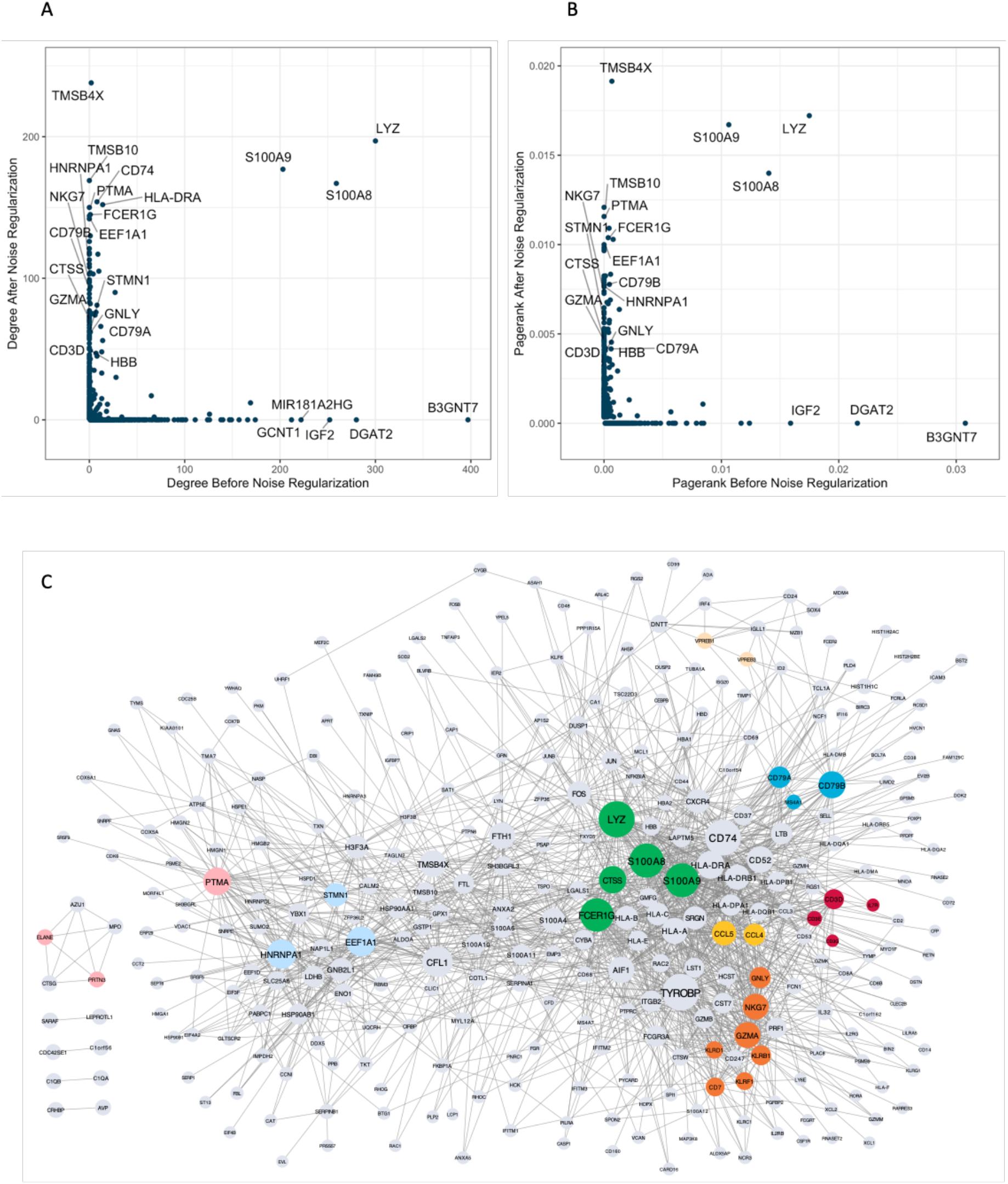
Gene-gene correlation network inferred from scRNA-seq data. (A) and (B) Compare degree and PageRank of each gene in the correlation networks constructed before and after noise regularization. Genes present in one network but not in the other were assigned a zero value in the non-presenting one. Selected genes with high degree/PageRank before or after noise regularization were labeled. Cell type marker genes such as NKG7, CD79B, HBB etc. had relatively higher degree and PageRank after noise regularization. (C) Network construction with refined gene-gene correlations (NBR + noise regularization + removing links not in PPI), the node size is proportional to its PageRank and the edge width is proportional to spearman correlation between the two genes (nodes). Cell type marker genes (colored nodes) such as CD79A, CD79B, NKG7, GNLY, LYZ, STMN1 have high PageRank, indicating their importance in different cell types. Cell type related genes also formed cell type specific modules.

## DISCUSSION

scRNA-seq technology has been gaining increasingly more popularity over the past decade. Proper and efficient data preprocessing are crucial for downstream analyses such as cell clustering, differential gene expression detection, and novel cell type discoveries^3,4^. Here, we benchmarked five data preprocessing methods for scRNA-seq with a focus on their influence in gene-gene correlation inference. Our results demonstrated that in a human bone marrow single cell dataset, all the methods except NormUMI generated inflated gene-gene correlations. Furthermore, the highly correlated gene pairs had low enrichment in PPI, indicating that they were more likely to be artifacts introduced during the data preprocessing procedure. Among these methods, NBR produced the lowest PPI enrichment, while NormUMI, method with the least data manipulation, yielded much higher enrichment as compared to the other four sophisticated methods. Thus, our benchmarking results raise the issue that correlation computed directly from these preprocessed data may not be reliable and should be treated with caution.

Manual inspection of the negative control results suggested that major causes of the spurious correlations may come from overfitting or over-smoothing during data preprocessing. The preprocessing methods, especially those imputing dropout events, rely heavily on internal similarity information (either gene-gene similarity or cell-cell similarity) within the original dataset. For instance, MAGIC uses the nearest neighbor graph to group cells with similar expression patterns and further imputes the missing values in the original data. Indeed, this could be circular to measure the gene-gene correlation after applying these steps. Given that these methods rely on the similarity of gene expression to amend gene expression, it is not surprising that they produce augmented gene-gene correlations.

To resolve the correlation artifact issues, we proposed a model-agnostic noise regularization method. False correlations from the overly smoothed data can be eliminated by the added noise while the true correlations should be robust enough to tolerate. Since the dynamic range of expression vary gene by gene, magnitude of the added noise should also be set relative to an individual gene’s expression level such that the true signal of genes with lower expression range can be preserved. Thus, the level of random noise is determined as a percentile of a gene’s dynamic range rather than a fixed value to be used for all genes. We further investigated the effect of different noise strengths (1, 5, 10, 20 percentile of the expression level), and found that the use of the 1 percentile produced the optimal PPI enrichment (**Fig. S3**). Finally, we generated random noise that ranged from 0 to 1 percentile of the gene expression level and applied them to the expression matrix. The noise regularization step remarkedly reduced the correlation artifacts and generated more reliable gene-gene association. However, it should be noted that the magnitude of the noise applied here was optimized to maximize the PPI enrichment, which may result in a higher true positive rate. Since there is always a trade-off between sensitivity and specificity, whether this noise strength is the optimal to reveal novel correlations likely requires further investigation.

Gene-gene correlations at the whole transcriptome level for bulk cells have been established to reconstruct gene-gene interaction networks and further uncover gene functions and genetic modules^22,28,29^. With the growing adoption of single cell technology, the use of scRNA-seq to infer gene-gene correlations and reconstruct global gene network is also burgeoning. Pioneering work by Iacono *et al*. used single-cell data-derived correlation metrics to generate gene regulatory networks and found that the networks could detect latent regulatory changes^30^. A deep learning approach has also been developed to predict transcription factor targets from single-cell expression data^31^. In this study, we used single cell gene-gene correlations derived after noise regularization to reconstruct a gene network that produced clear immune cell type related modules. We also evaluated the importance of each gene in the network by applying well established graph theory methods. We demonstrated that the canonical cell type markers yield higher degree and PageRank, in general, indicating their critical roles in different cell types.

A limitation of this study is that these methods were mostly implemented using their default parameters, which may not be optimal for this dataset. Changing the parameters and hyperparameters could have noticeable influence on the results. Andrews *et al*. tested different imputation methods on a simulation dataset and found that different parameters produced different degrees of false correlation^9^. Unfortunately, the choice of parameters is often arbitrary and lacks clear guidelines. For instance, MAGIC applies data smoothing by nearest neighbor graph. Increasing the number of neighbors will lead to smoother data, and in most cases resulting in inflated gene-gene correlations and higher false positives in correlation-based analyses. In addition, the diffusion time (*t*) in the algorithm also strongly affects the data smoothness. By default, this parameter is determined according to the Procrustes disparity of the diffused data. However, default setting apparently generated over-smoothed data in our study. Using different parameter value (e.g. decreased to a fixed number 6), we found that the output can be visually improved, but a high amount of spurious correlations still exists. This challenge is further complicated when users need to consider combinations of these parameters. A similar issue is also noticed in the implementation of DCA that requires a series of parameters, including many routine deep learning framework training parameters, such as learning rate and strength of L1/L2 regularization. The default architecture of DCA (three hidden layers with 64, 32, 64 neurons) was originally optimized on a simulation dataset with only 200 genes. When it is applied to real datasets that usually contain more than 10,000 genes, whether the default number of neurons can still capture the full picture and reconstruct reliable gene-gene networks becomes unclear. The lack of clear guidance for parameter selection could potentially hurt the reproducibility of any studies utilizing these preprocessing methods.

In summary, we compared five scRNA-seq data preprocessing methods on a real single-cell dataset and found that several preprocessing procedures may have introduced considerable amount of spurious gene-gene correlations. Therefore, single cell analysis involving gene-gene correlations should be performed with caution. To address the issue, we proposed a model-agnostic method to regularize the preprocessed data, which can effectively remove the spurious correlations and empower studies looking to reconstruct co-expression networks from scRNA-seq data.

## METHODS

### HCA scRNA-seq dataset

Bone marrow single-cell sequencing data was retrieved from Human Cell Atlas Data Portal (https://data.humancellatlas.org). The dataset contains profiling of 378,000 immunocytes by 10X Genomics chromium platform. Single-cell analysis was performed using Seurat R package (Version 3.0)^32^. In the quality control step, low quality cells were removed if they met one of the following criteria: (1) expressed less than 100 genes; (2) expressed more than 3,500 genes; (3) total UMI counts > 10,000; (4) mitochondrial RNA percentage > 10%. Remaining cells were clustered using k-nearest neighbor (KNN) graph-based clustering approach, with the first 30 principal components (PC) being used to construct the KNN graph. Clustering results were visualized with Uniform Manifold Approximation and Projection (UMAP), also using the first 30 PCs as inputs. In the subsequent correlation analysis, to reduce the computational burden, we randomly sampled 50,000 cells from the original dataset. We further filtered out genes expressed in less than 100 cells (0.2%), which left 12,600 genes remaining in the final benchmarking dataset.

### Normalization or imputation methods

NormUMI was performed using Seurat R package (Version 3.0) without log transformation^32^. NBR, SAVER and DCA were run with default parameters according to the software tutorials. Specifically, NBR was performed using sctransform R package (version 0.2.0)^33^. Poisson regression was performed for each gene under the negative binomial model. Regularized model parameters were used to transform observed UMI counts into Pearson residuals. DCA was performed with dca python package^13^: The deep learning framework had three hidden layers with 64, 32, 64 neurons. The learning rate used was 0.001 and batch size was set to 32. SAVER was run with SAVER R package (version 1.1.1) without requiring additional parameters^12^. MAGIC was run with MAGIC R implementation (version 1.5-9)^11^ with the following parameters: number of principle component npca= 30, the power of the Markov affinity matrix t=6 and number of nearest neighbor k= 30.

### Gene-gene correlation and mutual information calculation

Spearman correlation of each gene pair was calculated from cells in cluster 0 to cluster 9 (top 10 clusters with the largest cell number, which range from 583 to 16936 cells, **Fig. S1**), respectively. A gene was considered present in a cluster if its expression was detected in more than 1% of the cells or 50 cells in that cluster, whichever is greater. The correlation of a gene pair in one cluster was considered an effective correlation if the two genes were both considered as expressed in that cluster. The highest effective correlation across the ten clusters was recorded as the final correlation for a given gene pair. Mutual information of the selected gene pairs were measured using infotheo R package (version 1.2.0), data was discretized using the equal frequencies binning algorithm, and the entropy was estimated with an empirical probability distribution.

### Protein-protein interaction enrichment

Human protein-protein interaction data was retrieved from the STRING database (version 11) (http://string-db.org)^15^. The STRING database consists of comprehensively collected publicly available sources of protein–protein interaction information and is complemented with computational predictions. The final database includes both direct (physical) and indirect (functional) interactions. In version 11, 5,772,157 protein-protein interactions were included. After applying different data preprocessing methods, gene pairs were ranked by their spearman correlation coefficients. The top *n* gene pairs were then taken and overlapped with STRING database. The fraction of the top *n* pairs appearing in the database were recorded as the PPI enrichment.

### Noise regularization

Assuming that *V* is the expression value of gene *i* in cell *j* in the expression matrix processed by a specific method, a random noise value was generated and added to *V* by the following procedures: (1) Determine the expression distribution of gene *i* across all the cells. (2) Take 1 percentile of the gene *i* expression as the maximal noise level, denoted as *M*. If *M* equals to zero, 0.1 will be used as the maximal noise level. (3) Generate a random number ranging from 0 to *M* under uniform distribution and add this random number to *V* to get the noise regularized expression matrix.

### Network reconstruction

Within each cluster, we ranked the gene pairs by their spearman correlation coefficients. In this study, since we were more interested in cell type-specific gene interaction modules, we removed housekeeping genes from the network reconstruction. In general, housekeeping genes are required for basic cellular functions and thus expected to express regardless of cell types. The housekeeping gene list used here was obtained from a previous publication^34^, plus (i) typical housekeeping genes like ACTB, B2M, (ii) ribosomal, citrate cycle, and cytoskeleton genes from Reactome^35^ and (iii) mitochondrial DNA encoded genes. In total, 3,984 housekeeping genes were considered. After removing housekeeping genes, the top 1,000 gene pairs from each cluster were taken and put together to construct the draft network. The importance of each node in the network was measured by degree and PageRank using igraph R package^36^. We next refined the network by removing links that do not overlap with PPI in STRING database. The final network was visualized using Cytoscape ^37^ together with R package RCy3^38^. The network layout was generated using EntOptLayout Cytoscape plug-in^27^.

## Code availability

The R code for analyses in this study will be available at Github: https://github.com/RuoyuZhang/NoiseRegularization.

## Acknowledgements

We thank human cell atlas for generating “Census of Immune Cells” dataset and make it available to the research community. We thank Drs. Ian Setliff and Kaitlyn Gayvert for their helpful discussion and comments on the manuscript.

---

**Figure S1.**
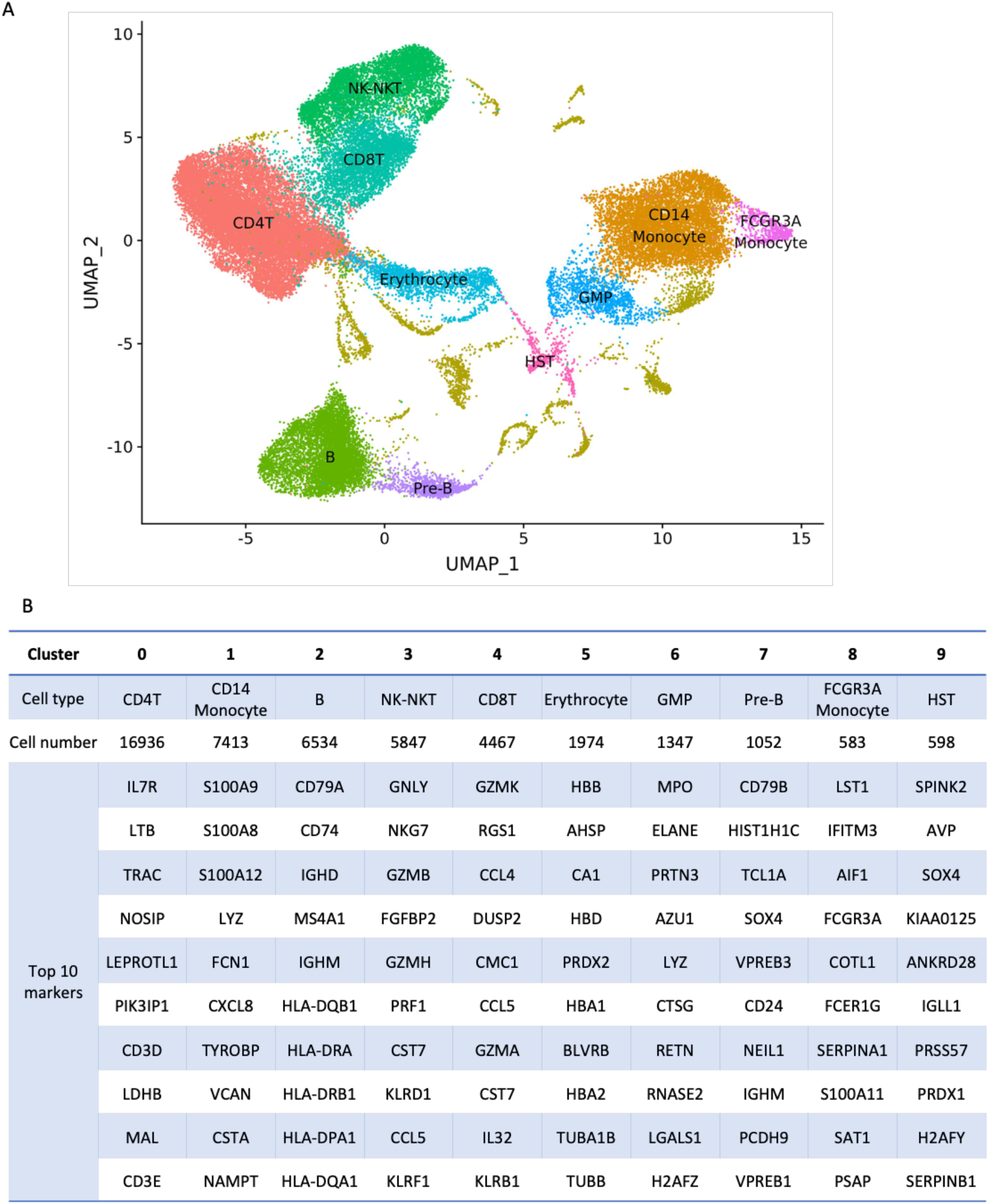
HCA single cell data used for this study. (A) UMAP of 50,000 bone marrow cells, covering major immune cell types. (B) Cell type annotations, cell counts and the top 10 markers for the 10 biggest clusters in this dataset.

**Figure S2.**
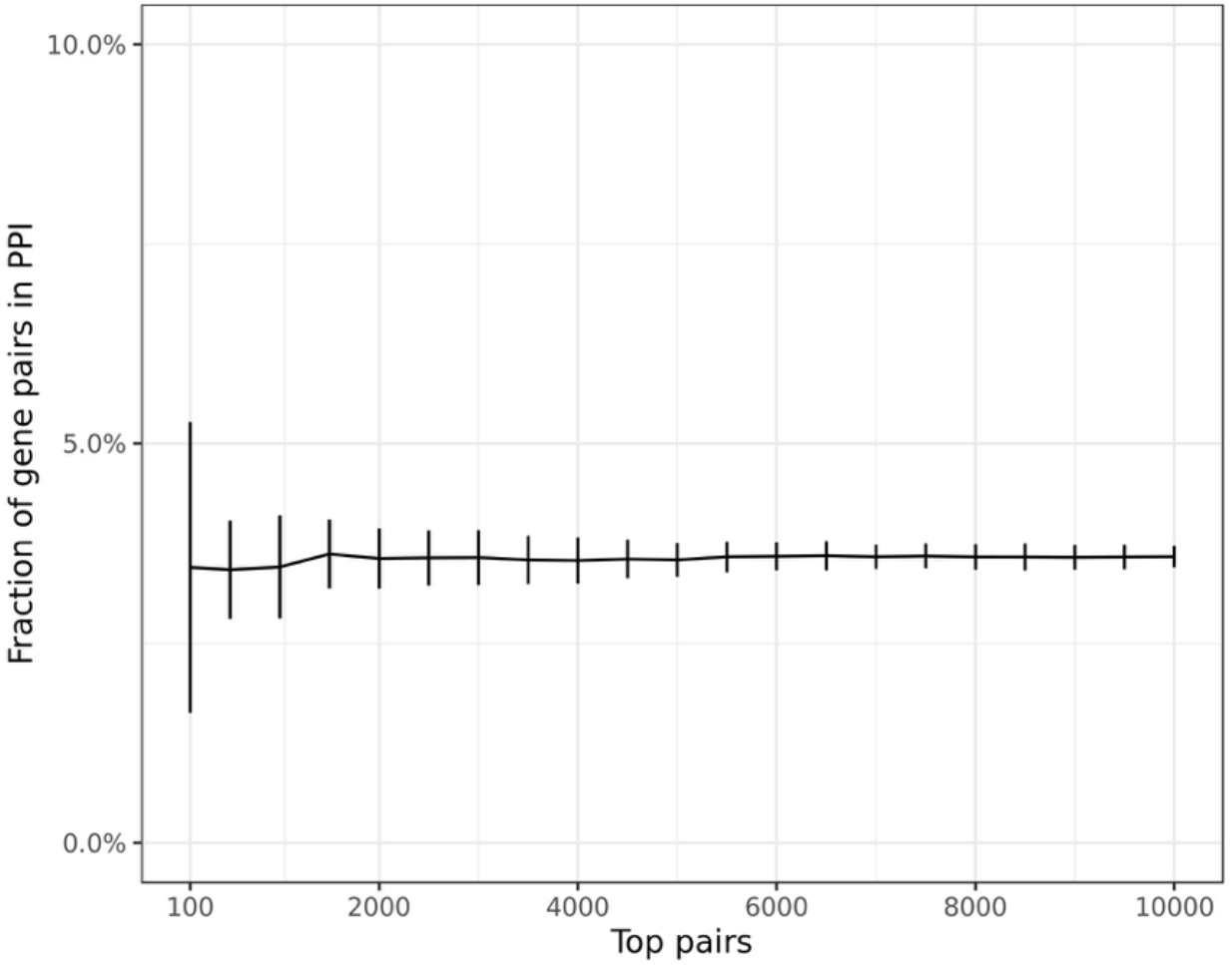
PPI enrichment of randomly sampled gene pairs. Gene pairs were randomly sampled and overlapped with PPI database to estimate the background enrichment level. The mean of the background enrichment is ~3.6%, error bar represents one standard deviation based on 20 random samplings.

**Figure S3.**
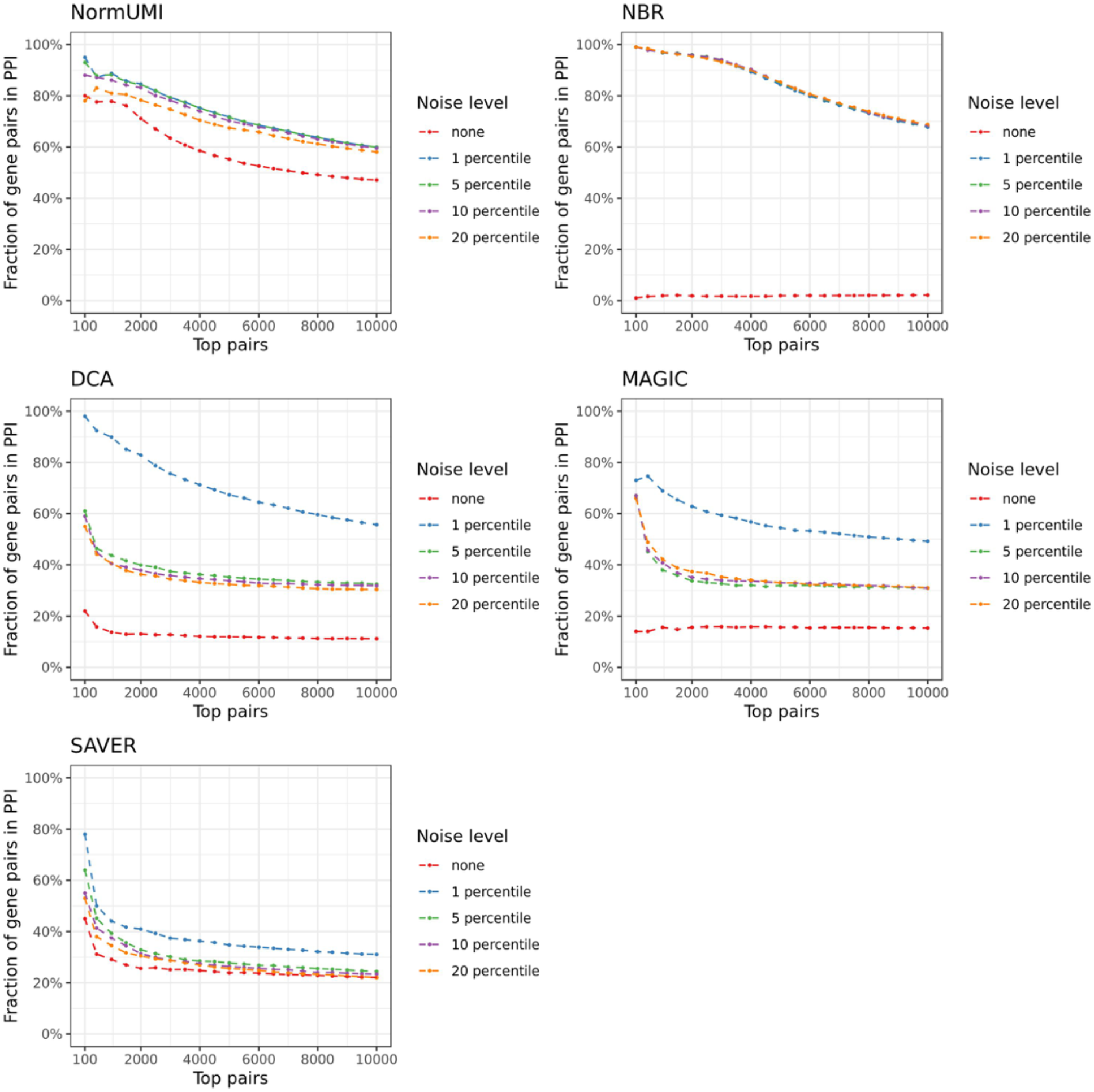
PPI enrichment after adding noise at different levels. Different level of noise is applied to regularize the data (1, 5, 10, 20 percentile of the expression level). Noise at 1 percentile of the expression level produces the optimal PPI enrichment.

## Reference

1 Freeman, T. C. et al. Construction, visualisation, and clustering of transcription networks from microarray expression data. PLoS computational biology 3, e206.

2 Ballouz, S., Verleyen, W. & Gillis, J. Guidance for RNA-seq co-expression network construction and analysis: safety in numbers. Bioinformatics 31, 2123–2130.

3 Kolodziejczyk, Aleksandra A., Kim, J. K., Svensson, V., Marioni, John C. & Teichmann, Sarah A. The Technology and Biology of Single-Cell RNA Sequencing. Molecular Cell 58, 610–620.

4 Papalexi, E. & Satija, R. Single-cell RNA sequencing to explore immune cell heterogeneity. Nature Reviews Immunology 18, 35.

5 Hicks, S. C., Townes, F. W., Teng, M. & Irizarry, R. A. Missing data and technical variability in single-cell RNA-sequencing experiments. Biostatistics 19, 562–578.

6 Svensson, V. et al. Power analysis of single-cell RNA-sequencing experiments. Nature Methods 14, 381.

7 Ziegenhain, C. et al. Comparative Analysis of Single-Cell RNA Sequencing Methods. Molecular Cell 65, 631–643.e634.

8 Tian, L. et al. Benchmarking single cell RNA-sequencing analysis pipelines using mixture control experiments. Nature Methods 16, 479–487.

9 Andrews, T. & Hemberg, M. False signals induced by single-cell imputation [version 1; peer review: 4 approved with reservations]. F1000Research 7.

10 Hafemeister, C. & Satija, R. Normalization and variance stabilization of single-cell RNA-seq data using regularized negative binomial regression. Genome Biology 20, 296.

11 van Dijk, D. et al. Recovering Gene Interactions from Single-Cell Data Using Data Diffusion. Cell 174, 716–729.e727.

12 Huang, M. et al. SAVER: gene expression recovery for single-cell RNA sequencing. Nature Methods 15, 539–542.

13 Eraslan, G., Simon, L. M., Mircea, M., Mueller, N. S. & Theis, F. J. Single-cell RNA-seq denoising using a deep count autoencoder. Nature Communications 10, 390.

14 Regev, A. et al. The Human Cell Atlas. eLife 6, e27041.

15 Szklarczyk, D. et al. STRING v11: protein–protein association networks with increased coverage, supporting functional discovery in genome-wide experimental datasets. Nucleic Acids Research 47, D607–D613.

16 Bishop, C. M. Training with noise is equivalent to Tikhonov regularization. Neural computation 7, 108–116.

17 Neelakantan, A. et al. Adding gradient noise improves learning for very deep networks. arXiv preprint arXiv:1511.06807.

18 Smilkov, D., Thorat, N., Kim, B., Viégas, F. & Wattenberg, M. Smoothgrad: removing noise by adding noise. arXiv preprint arXiv:1706.03825.

19 Ashburner, M. et al. Gene ontology: tool for the unification of biology. The Gene Ontology Consortium. Nature genetics 25, 25–29.

20 The Gene Ontology Consortium. The Gene Ontology Resource: 20 years and still GOing strong. Nucleic Acids Research 47, D330–D338.

21 Kinney, J. B. & Atwal, G. S. Equitability, mutual information, and the maximal information coefficient. Proceedings of the National Academy of Sciences 111, 3354–3359.

22 Stuart, J. M., Segal, E., Koller, D. & Kim, S. K. A Gene-Coexpression Network for Global Discovery of Conserved Genetic Modules. Science 302, 249–255.

23 Bondy, J. A. & Murty, U. S. R. Graph Theory. (Springer Publishing Company, Incorporated, 2008).

24 Page, L., Brin, S., Motwani, R. & Winograd, T. The PageRank citation ranking: Bringing order to the web. (Stanford InfoLab, 1999).

25 Cheng, H., Jiang, L., Wu, M. & Liu, Q. Inferring Transcriptional Interactions by the Optimal Integration of ChIP-chip and Knock-out Data. Bioinform Biol Insights 3, 129–140.

26 Sayyed-Ahmad, A., Tuncay, K. & Ortoleva, P. J. Transcriptional regulatory network refinement and quantification through kinetic modeling, gene expression microarray data and information theory. BMC Bioinformatics 8, 20.

27 Ágg, B. et al. The EntOptLayout Cytoscape plug-in for the efficient visualization of major protein complexes in protein–protein interaction and signalling networks. Bioinformatics.

28 Costanzo, M. et al. The Genetic Landscape of a Cell. Science 327, 425–431.

29 Carro, M. S. et al. The transcriptional network for mesenchymal transformation of brain tumours. Nature 463, 318–325.

30 Iacono, G., Massoni-Badosa, R. & Heyn, H. Single-cell transcriptomics unveils gene regulatory network plasticity. Genome Biology 20, 110.

31 Yuan, Y. & Bar-Joseph, Z. Deep learning for inferring gene relationships from single-cell expression data. Proceedings of the National Academy of Sciences 116, 27151–27158.

32 Butler, A., Hoffman, P., Smibert, P., Papalexi, E. & Satija, R. Integrating single-cell transcriptomic data across different conditions, technologies, and species. Nature Biotechnology 36, 411.

33 Hafemeister, C. & Satija, R. Normalization and variance stabilization of single-cell RNA-seq data using regularized negative binomial regression. bioRxiv, 576827.

34 Eisenberg, E. & Levanon, E. Y. Human housekeeping genes, revisited. Trends in Genetics 29, 569–574.

35 Fabregat, A. et al. The Reactome Pathway Knowledgebase. Nucleic Acids Research 46, D649–D655.

36 Csardi, G. & Nepusz, T. The igraph software package for complex network research. InterJournal, Complex Systems 1695, 1–9.

37 Shannon, P. et al. Cytoscape: A Software Environment for Integrated Models of Biomolecular Interaction Networks. Genome Research 13, 2498–2504.

38 Ono, K., Muetze, T., Kolishovski, G., Shannon, P. & Demchak, B. CyREST: Turbocharging Cytoscape Access for External Tools via a RESTful API. F1000Research 4, 478–478.

